# Cholesterol metabolism drives regulatory B cell function

**DOI:** 10.1101/2020.01.03.893982

**Authors:** Jack A. Bibby, Harriet A. Purvis, Thomas Hayday, Anita Chandra, Klaus Okkenhaug, Michael Wood, Helen J. Lachmann, Claudia Kemper, Andrew P. Cope, Esperanza Perucha

**Author notes:** These authors contributed equally to this work (Andrew P. Cope, Esperanza Perucha).

## Abstract

Regulatory B cells restrict immune and inflammatory responses across a number of contexts. This capacity is mediated primarily through the production of IL-10. Here we demonstrate that the induction of a regulatory program in human B cells is dependent on a metabolic priming event driven by cholesterol metabolism. Synthesis of the metabolic intermediate geranylgeranyl pyrophosphate (GGPP) was required to specifically drive IL-10 production, and to attenuate Th1 responses. Furthermore, GGPP-dependent protein modifications controlled signaling through PI3Kδ-AKT-GSK3, which in turn promoted BLIMP1-dependent IL-10 production. Inherited gene mutations in cholesterol metabolism result in a severe autoinflammatory syndrome, termed mevalonate kinase deficiency (MKD). Consistent with our findings, B cells from MKD patients induced poor IL-10 responses and were functionally impaired. Moreover, metabolic supplementation with GGPP was able to reverse this defect. Collectively, our data define cholesterol metabolism as an integral metabolic pathway for the optimal functioning of human IL-10 producing regulatory B cells.

**Graphical abstract:** MKD = Mevalonate kinase deficiency

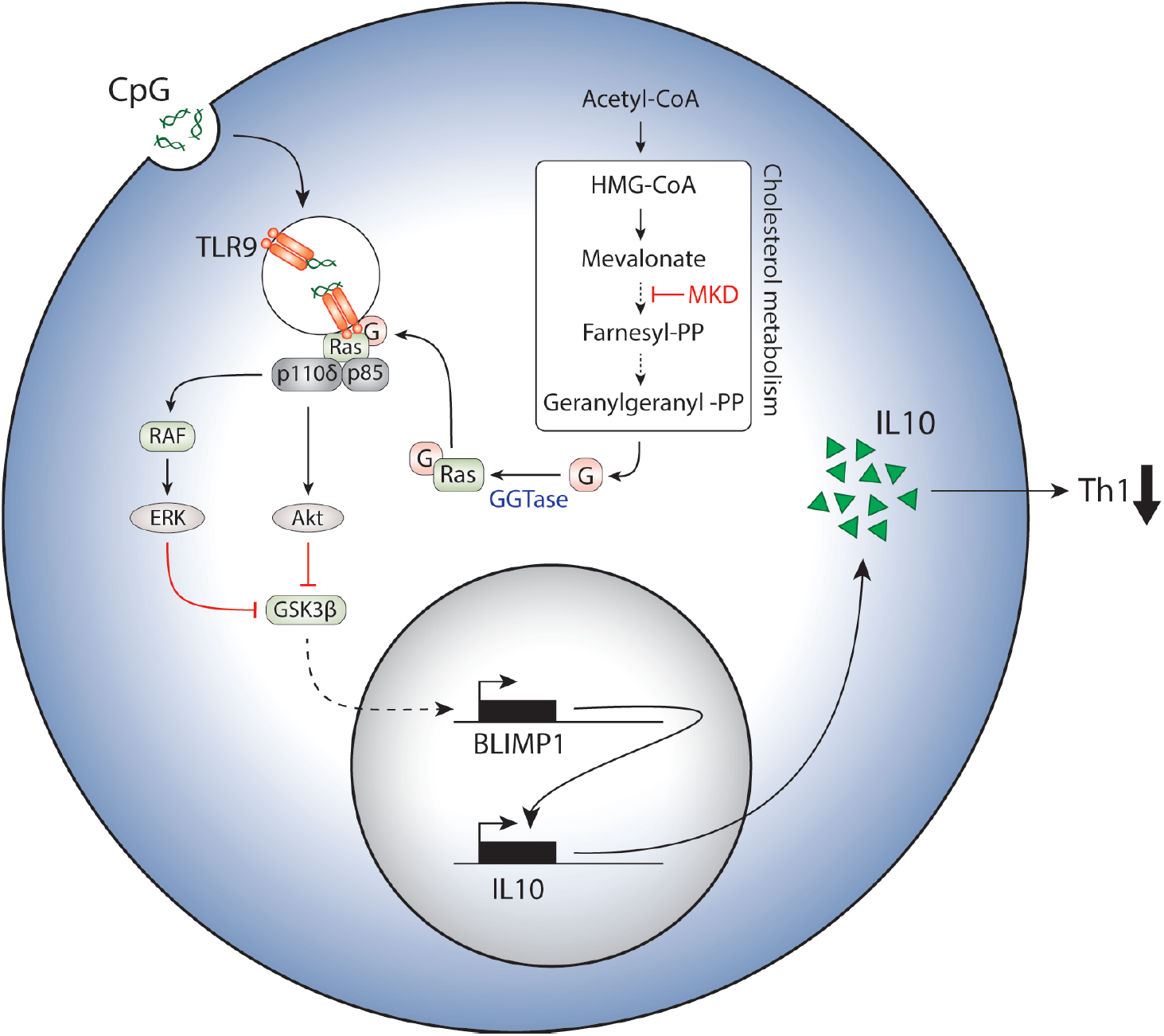

## Introduction

Immunosuppressive B cells form a critical component of the immune regulatory compartment ^1,2^. It is thought that their suppressive capacity derives mainly from their ability to produce IL-10, and in the absence of any lineage marker, this is considered a hallmark of regulatory B cells ^3–6^. Their functional importance has been well described in murine models of disease, demonstrating a potent regulatory capacity across a number of contexts including infection, cancer, and autoimmune disease ^3,7–10^. Comparable suppressive activity has been demonstrated in human regulatory B cells *in vitro*, suggesting that these cells also contribute toward regulating inflammatory responses in humans ^5,6^. In support of this, IL-10 producing B cells have been demonstrated to be numerically depleted or functional impaired, *ex vivo*, in patients with inflammatory disease ^5,11,12^.

Despite the importance of IL-10 production by B cells, relatively little is known about the molecular mechanisms that govern its expression. Typically, induction of a regulatory phenotype in both human and murine B cells has been achieved through ligation of TLR9 or CD40 ^5,6,13^. Downstream, PI3K and ERK signals appear to be important for IL-10 expression ^14,15^. This is in broad agreement with *in vivo* data from mouse models suggesting that both TLR and CD40 signals are required for the induction of IL-10 in response to inflammatory stimuli ^3,13^. However, precise details of the signaling cascades or cellular profiles underpinning the induction of a regulatory program in B cells are poorly understood.

Control of immune cell metabolism is critical in regulating fundamental immunological processes^16,17^. However, in comparison to innate cell and T cell lineages, there has been relatively little work aimed at understanding the metabolic regulation of B cell biology ^18–20^. Furthermore, there is currently no understanding toward the metabolic requirements of regulatory B cells. An emerging concept from studies of regulatory T cells and M2 macrophages is their heightened reliance on lipid metabolism in comparison to the metabolic requirements of inflammatory immune cell lineages. Much of this work has focused on fatty acid oxidation, defining this as a central pathway that underpins polarization and effector functions in regulatory cells ^21^. In contrast, the contribution of cholesterol metabolism (the multi-step conversion of HMG-CoA to cholesterol) to immune function is less well characterized. Our recent studies, as well as those of others, suggest that cholesterol metabolism plays a role in restricting inflammatory responses ^22–24^. These data are consistent with patients carrying mutations in the pathway, who develop severe and recurring autoinflammatory fevers, associated with dysregulated B cell responses, manifest by elevated serum immunoglobulin levels ^25^.

Given the immunoregulatory associations with cholesterol metabolism, we set out to address its role in IL-10 producing B cells. In doing so, we revealed that cholesterol metabolism is a central metabolic pathway required for the induction of a regulatory phenotype in human B cells, defining the specific metabolic intermediate of the pathway that mediates critical signaling events for the induction of a regulatory B cell program. In vivo evidence for this specific metabolic requirement is provided, by virtue of defective regulatory responses of B cells derived from patients with inherited defects in cholesterol metabolism.

## Results

### Cholesterol metabolism drives regulatory B cell function via IL-10

Given that patients with dysregulated cholesterol metabolism generate severe systemic inflammation and hyperactive B cell responses, we investigated the role of cholesterol metabolism in regulatory B cell function (the metabolic pathway is depicted in Supplementary Figure 1). Consistent with previous observations, TLR9 ligation in human B cells induces a potent IL-10 response (Figure 1A-B), which mediates their regulatory phenotype ^5,11^. In agreement, TLR9 activated B cells were able to suppress Th1 induction in CD4^+^ T cells *in vitro* (protocol outlined in Figure 1C, Figure 1D). Neutralization of IL-10 resulted in a complete loss of suppression, confirming the dependence on IL-10 in mediating this function (Figure 1D). In line with this, concentrations of IL-10 produced by B cells upon TLR9 stimulation (>1ng/ml, Figure 1B) were sufficient to directly suppress CD4^+^ T cell IFNγ expression (Supplementary Figure 2). In contrast, pre-treatment of B cells with the HMG-CoA reductase inhibitor atorvastatin, which inhibits an early step of the cholesterol metabolic pathway, abrogated this suppressive capacity. Furthermore, supplementation of exogenous mevalonate reversed this defect, suggesting the requirement for a specific metabolite downstream of HMG-CoA (Figure 1D).

**Figure. 1.**
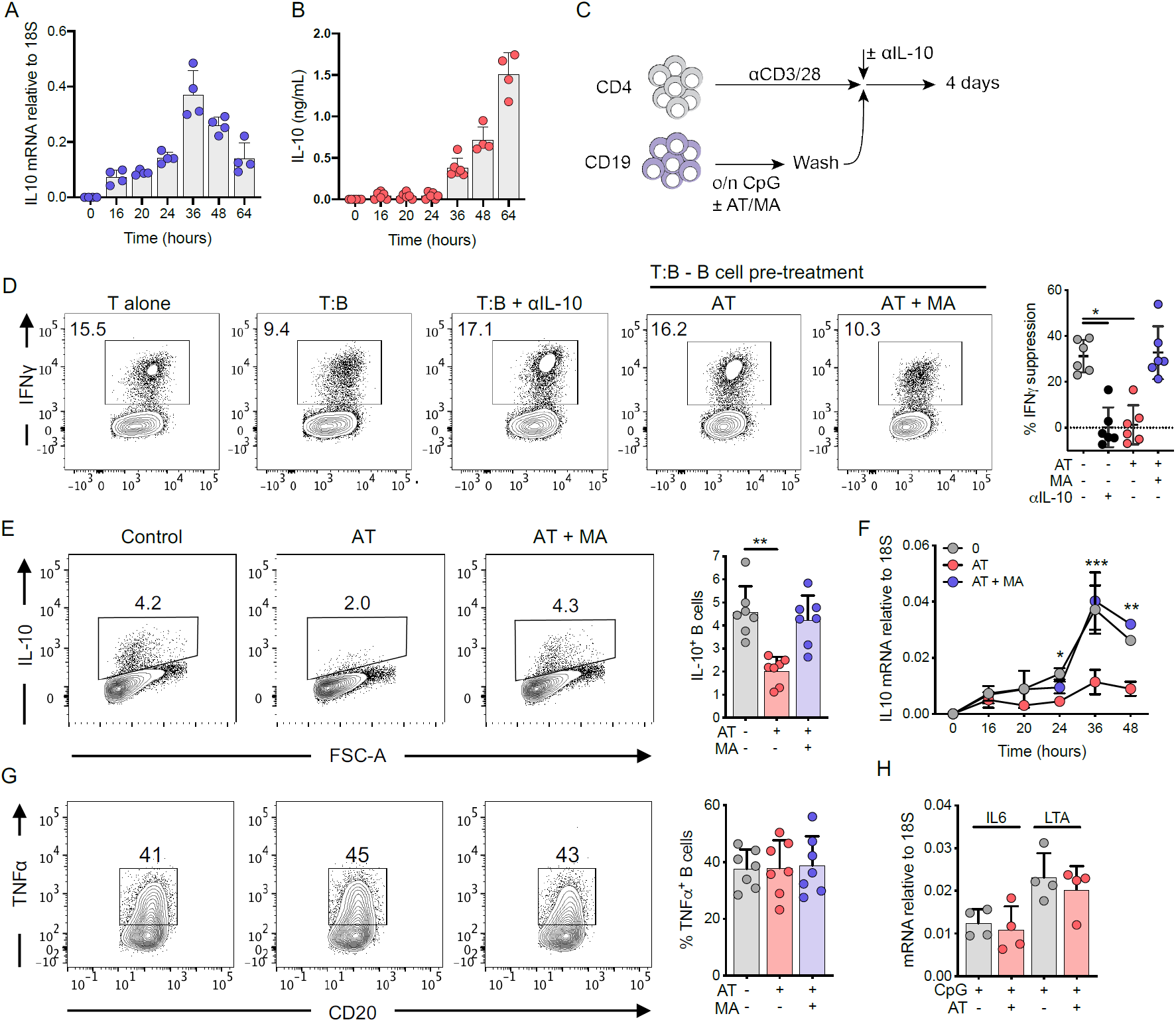
Cholesterol metabolism drives regulatory B cell function via IL-10. **A-B.** Kinetics of IL-10 mRNA transcript **(A)** and protein **(B)** expression at various time points in human B cells after TLR9 stimulation (n=4 for mRNA or n=4-5 for protein). IL10 mRNA was measured by qRT-PCR, and calculated relative to 18S by the formula 2–ΔCt. **C.** Schematic protocol for the co-culture for T and B cells. Briefly, human CD19^+^ B cells were stimulated with CpG ± atorvastatin (AT) ± mevalonate (MA) overnight, before washing and addition to αCD3/28 stimulated human CD4^+^ T cells for 4 days ± αIL-10 antibody. **D.** IFNγ production in human CD4^+^ T cells after co-culture with autologous TLR9-activated B cells. IFNγ suppression was calculated by: ((x-y)/x)*100 where x = IFNγ production by T cells alone, y = IFNγ production in co-cultured T cells. **E.** IL-10 expression in human B cells after stimulation through TLR9 ± AT ± MA. **F.** IL-10 mRNA expression over time in human B cells after stimulation through TLR9 ± AT ± MA. **G.** TNFα expression in human B cells after stimulation through TLR9 ± AT ± MA. **H.** IL6 or LTA mRNA expression, relative to 18S, over time in human B cells after stimulation through TLR9 ± AT. Each data point represents individual donors. All data presented are mean ± SD. \**P*<0.05, \*\**P*<0.01, \*\*\**P*<0.001, and all significant values are shown. Statistical analysis was conducted using Friedman’s test with Dunn’s multiple comparisons

The loss of suppressive capacity by IL-10 neutralization and HMG-CoA reductase inhibition indicated that cholesterol metabolism could be directly regulating IL-10 production. Indeed, inhibition of HMG-CoA reductase impaired the ability of B cells to produce IL-10, both at the mRNA and protein level (Figure 1E-F, Supplementary Figure 3A-B). Once again, and consistent with metabolic depletion downstream of HMG-CoA, exogenous mevalonate was able to reverse this defect (Figure 1E-F). In contrast to IL-10, we observed a concurrent preservation in the inflammatory response, as determined by TNFα and IFNγ protein, and IL6, and LTA gene expression (Figure 1G-H, Supplementary Figure 3C-D). Together, these data indicated that cholesterol metabolism was critical in mediating IL-10 expression, and therefore the anti-inflammatory function of human B cells.

### Cholesterol metabolism drives IL-10 independent of B cell population

We next aimed to understand how cholesterol metabolism was able to mediate IL-10 production. Certain populations of human B cells have been proposed as primary producers of IL-10. The most well characterized of these are CD24^hi^CD27^+^ (B10) and CD24^hi^CD38^hi^ B cells ^5,6^. In agreement with previous observations, tSNE analysis demonstrated that all populations measured (B10, CD24^hi^CD38^hi^, naïve, memory, and plasmablast) contribute to the pool of IL-10 expressing cells to varying degrees (IL-10^+^ cells ranging from 1-12% of B cell populations, Supplementary Figure 4A). Furthermore, B10 and CD24^hi^CD38^hi^ B cells produced higher levels (2-3-fold) of IL-10 in response to TLR9 stimulation (Supplementary Figure 4B-C). Following inhibition of HMG-CoA reductase we observed no change in frequencies of B cell populations, viability, or cell surface markers (HLA-DR, CD86, or CD40), excluding the possibility that perturbation of cholesterol metabolism was depleting specific B cell subsets that possess a greater ability to express IL-10 (Supplementary Figure 4D-F). Furthermore, HMG-CoA reductase inhibition resulted in a 2-3-fold reduction in IL-10 expression irrespective of phenotype (Supplementary Figure 4G). Therefore, these data indicated a role for cholesterol metabolism in regulating IL-10 production that is shared across B cell populations, rather than an effect on specific populations.

### Cholesterol metabolism drives IL-10 via geranylgeranyl pyrophosphate

To more precisely understand the mechanistic control by cholesterol metabolism, we next sought to investigate if a specific pathway metabolite downstream of HMG-CoA was regulating IL-10. Cholesterol metabolism encompasses a number of metabolic pathways implicated in immune function including mevalonate, isoprenyl and sterol metabolism (Supplementary Figure 1), all of which are attenuated by HMG-CoA reductase inhibition to varying degrees. Given that defects in the isoprenyl branch have been demonstrated to result in hyperinflammatory responses *in vivo* ^23,26^, we investigated if isoprenylation was regulating IL-10. To this end, we targeted geranylgeranyltransferase (GGTase) and farnesyltransferase (FTase), enzymes known to post-translationally modify proteins with geranylgeranyl pyrophosphate (GGPP) or farnesyl pyrophosphate (FPP) groups respectively (enzymes and metabolites outlined in Figure 2A). Inhibition of GGTase, but not FTase, prevented TLR9-induced IL-10 production, indicating that geranylgeranyl dependent modifications regulate IL-10 expression (Figure 2B, Supplementary Figure 5A). In keeping with the effects of HMG-CoA reductase inhibition, inflammatory cytokine production was preserved (Figure 2C). To support the notion of GGPP dependency, we depleted metabolites within the pathway with atorvastatin, and found that specifically supplementing the geranyl branch with exogenous GGPP prevented the blockade of IL-10 production (Figure 2D, Fig S5B). Finally, to understand if cellular utilization of GGPP was dependent on the enzymatic activity of GGTase, we supplied cells with exogenous GGPP together with inhibition of GGTase. We found that GGPP was unable to rescue levels of IL-10 (Figure 2E), consistent with the idea that both GGTase activity and GGPP sufficiency are required for TLR9 induced expression of IL-10.

**Figure. 2.**
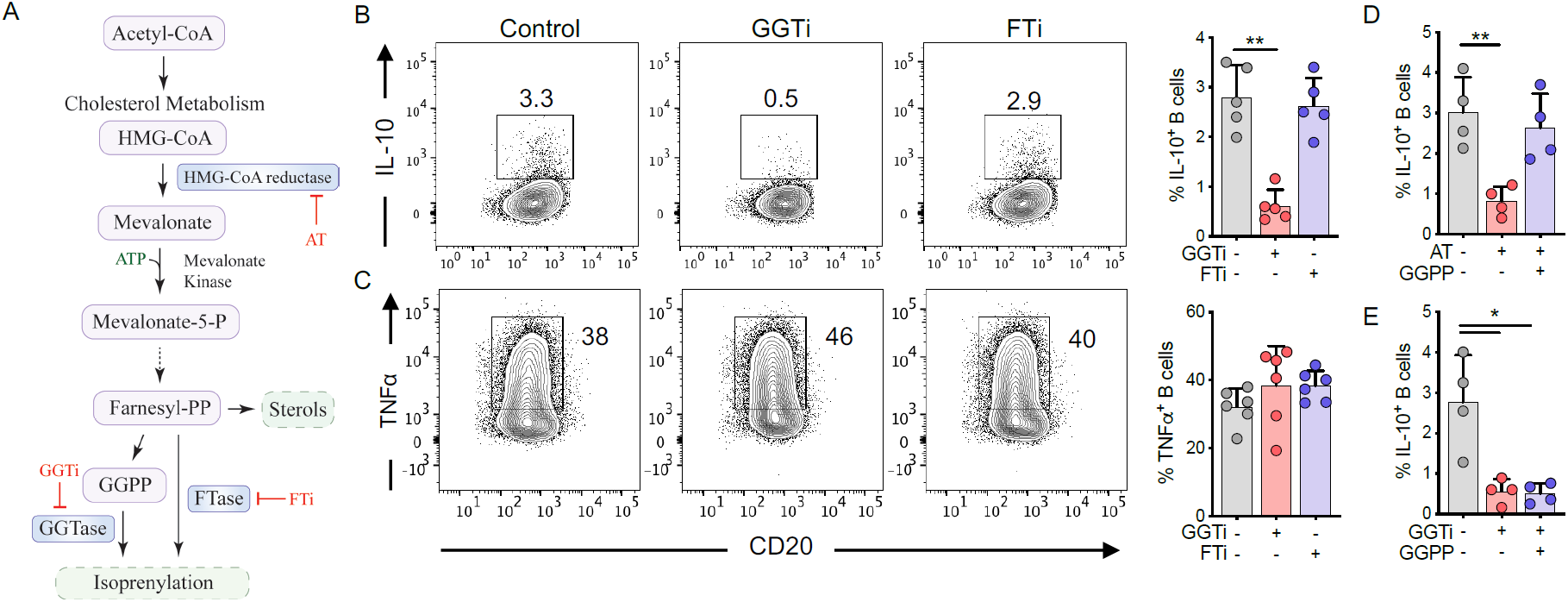
Cholesterol metabolism drives IL-10 via geranylgeranyl pyrophosphate. **A.** Schematic diagram showing key metabolites and enzymes of the isoprenylation route in cholesterol metabolism. **B-C.** IL-10 **(B)** and TNFα **(C)** expression in human B cells after stimulation through TLR9 ± geranylgeranyl transferase inhibition (GGTi) ± farnesyl transferase inhibition (FTi). **D.** IL-10 expression in human B cells after stimulation through TLR9 ± atorvastatin (AT) ± geranylgeranyl pyrophosphate (GGPP). **E.** IL-10 expression in human B cells after stimulation through TLR9 ± GGTi ± GGPP. Each data point represents individual donors. All data presented are mean ± SD. \**P*<0.05, \*\**P*<0.01, and all significant values are shown. Statistical analysis was conducted using Friedman’s test with Dunn’s multiple comparisons

### Geranylgeranyl pyrophosphate regulates signaling through TLR9

Isoprenyl modifications almost exclusively regulate the localization of Ras superfamily proteins ^27^. Ras-dependent pathways downstream of TLR9 include Raf-MEK-ERK and PI3K-AKT cascades (Figure 3A). Signaling through both pathways is critically required for IL-10 production, as inhibition of either pathway is sufficient to block TLR9-dependent IL-10 induction (Figure 3B-C). Therefore, we sought to address if activation of these pathways is dependent on GGPP. Following GGTase inhibition, AKT phosphorylation on Ser473 was severely impaired, whereas ERK phosphorylation was modestly reduced at early timepoints (Figure 3D, quantification shown in Supplementary Figure 6A-B). Downstream, AKT restricts GSK3 activity through an inhibitory phosphorylation event targeting Ser9 ^28^. In keeping with the idea that GSK3 suppression is required for IL-10 production, chemical inhibition of GSK3 enhanced TLR9-induced IL-10 expression (Figure 3E). Blocking GGTase activity resulted in reduced Ser9 phosphorylation on GSK3. This indicated preserved activation of GSK3, and that GGTase activity negatively regulates GSK3, which in turn is necessary for IL-10 production (Figure 3F). We then examined if inhibition of GSK3 was sufficient to rescue an upstream perturbation of geranylgeranylation. Indeed, bypassing GGTase through GSK3 inhibition was able to fully rescue IL-10 expression, without affecting TNFα (Figure 3G). Together these data suggest that following TLR9 engagement, IL-10 induction through AKT-GSK3, and possibly ERK, is dependent on GGTase activity.

**Figure. 3.**
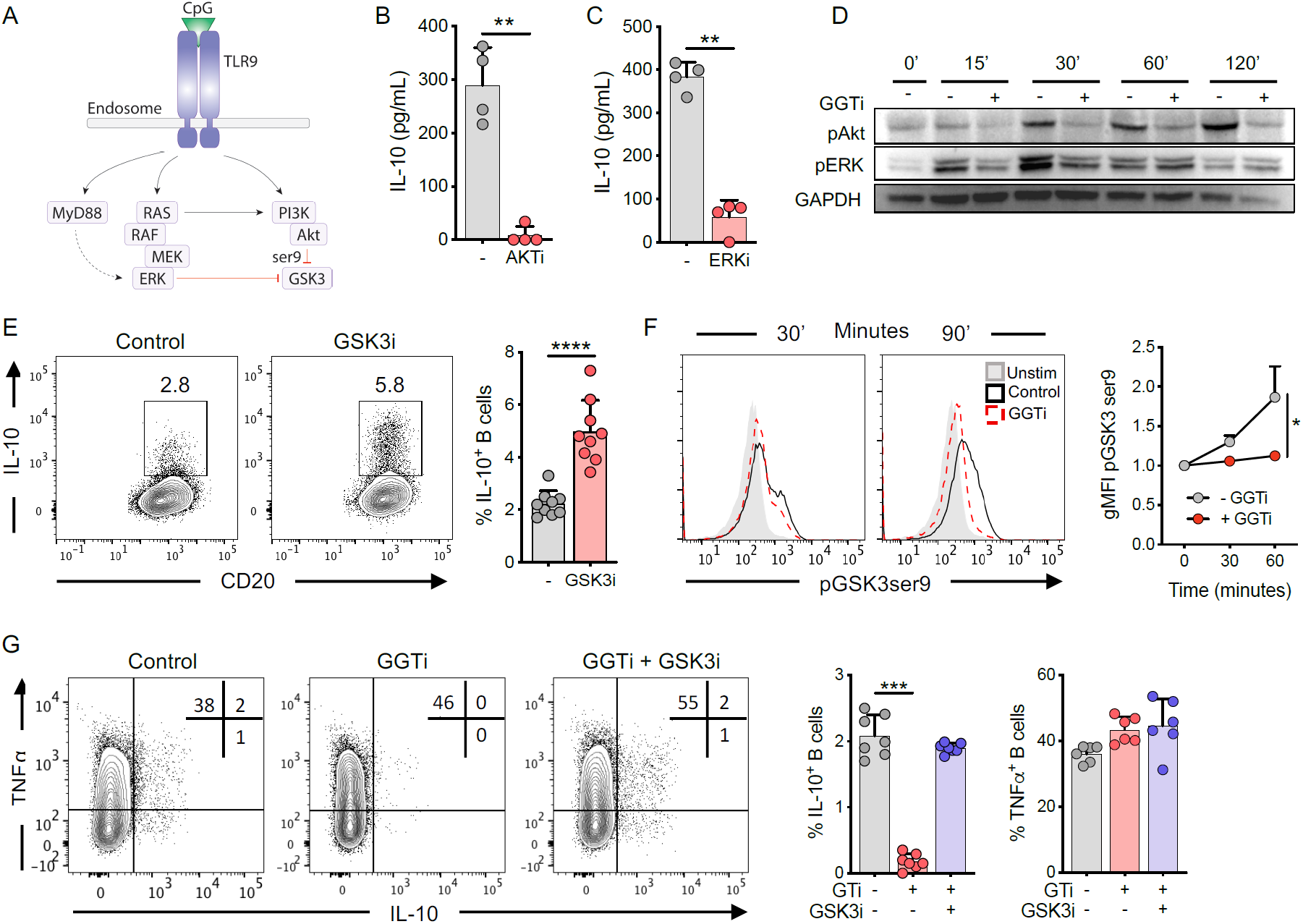
Geranylgeranyl pyrophosphate regulates signalling through TLR9. **A.** Schematic of Ras-dependent signalling cascades downstream of TLR9. **B-C.** IL-10 expression in human CD19^+^ B cells stimulated through TLR9 ± inhibition of **(B)** AKT (AKTi) or **(C)** ERK (ERKi). **D.** Phosphorylation of AKT (ser473) or ERK1/2 (Thr202/Tyr204) over time in human B cells upon stimulation through TLR9 ± inhibition of geranylgeranyl transferase (GGTi). **E.** IL-10 expression in human B cells stimulated through TLR9 ± inhibition of GSK3 (GSK3i). **F.** Phosphorylation of GSK3 (ser9) over time in human B cells upon stimulation through TLR9 ± inhibition of geranylgeranyl transferase (GGTi), relative to unstimulated controls. **G.** IL-10 and TNFα expression in human B cells stimulation through TLR9 ± GGTi ± GSK3i. Each data point represents individual donors. All data presented are mean ± SD. \*\**P*<0.01, \*\*\**P*<0.001, \*\*\*\**P*<0.0001 and all significant values are shown. Statistical analysis was conducted using Friedman’s test with Dunn’s multiple comparisons, or a paired t-test

### PI3Kδ regulates IL-10 expression in human and murine B cells

We next aimed to determine how geranylgeranyl-driven phosphorylation cascades regulate AKT-GSK3 signaling. PI3K mediates Ras-dependent AKT signaling, which suggests that isoprenylation-driven phosphorylation cascades through AKT are dependent on PI3K activity. Accordingly, pan inhibition of PI3K blocked expression of IL-10 upon TLR9 stimulation (Figure 4A). We found, by selective inhibition of either δ, α, or γ isoforms of PI3K, that IL-10 is primarily regulated through PI3Kδ downstream of TLR9 (Figure 4B). To further support the importance of PI3Kδ, we examined IL-10 production in splenic B cells derived from mice expressing either a hyperactive (E1020K) or catalytically inactive (D910A) PI3Kδ subunit. Following TLR9 activation, presence of the hyperactive PI3Kδ mutant resulted in 3-fold increased expression of IL-10, whereas the catalytically inactive PI3Kδ resulted in almost a complete loss of IL-10 production (Figure 4C). Furthermore, and in agreement with our previous observations, TNFα production was preserved (Figure 4C). These data suggest that isoprenylation-dependent interactions between Ras and PI3Kδ are required for IL-10 production, and likely underpin the regulatory function of B cells.

**Figure. 4.**
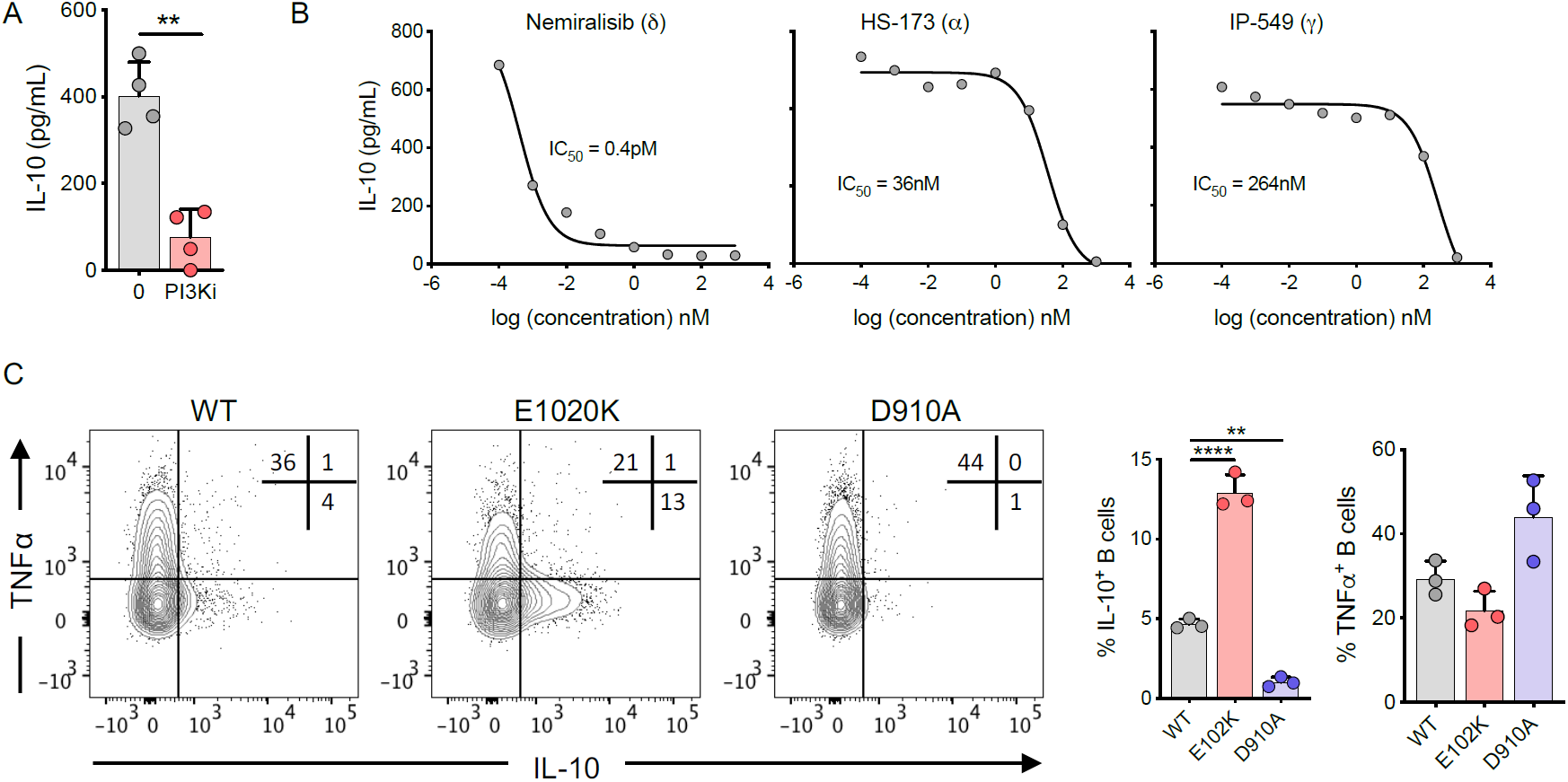
PI3Kδ regulates IL-10 expression in human and murine B cells. **A.** IL-10 expression in human B cells stimulated through TLR9 ± pan inhibition of PI3K (PI3Ki, inhibitor used was ZSTK474). **B.** IL-10 expression in human B cells stimulated through TLR9 ± inhibitors for specific family members of PI3K: these being δ (Nemiralisib), γ (IP-549), or α (HS-173), over a range of concentrations. N=4. **C.** IL-10 and TNFα expression in murine B cells after stimulation through TLR9. Mice used were wild type (WT), hyperactive PI3Kδ (E1020K), or catalytically inactive PI3Kδ (D910A). Each data point represents individual donors or mice. Data presented are mean ± SD. \*\**P*<0.01, \*\*\*\**P*<0.0001 and all significant values are shown. Statistical analysis was conducted using Friedman’s test with Dunn’s multiple comparisons (C), or a paired t-test (A)

### Geranylgeranyl pyrophosphate regulates IL-10 induction via BLIMP1

Currently there is no defined transcription factor that regulates IL-10 in human B cells. Therefore, we sought to understand how IL-10 is transcriptionally regulated, and to clarify the role for GGPP in this process. Stimulation of B cells in the presence of actinomycin D indicated that a transcriptional event within the first 24 hours was necessary for IL-10 production (Supplementary Figure 7A). Therefore, we suspected that under conditions of GGTase inhibition, expression of a transcription factor necessary for IL-10 may be compromised. To test this, we performed RNA-sequencing on TLR9-stimulated human B cells in the presence or absence of GGTase inhibition. Principal component analysis demonstrated that the main variation in the transcriptional profile of the cells was dependent on GGTase activity, as expected (Figure 5A, Supplementary Figure 7B). We then interrogated the list of differentially expressed genes (±1.5 fold, FDR<0.05) for those known or likely to encode human transcription factors (as defined by ^29^). Among the 73 differentially expressed transcription factors, 9 have been shown to regulate IL-10 expression in other cell types (Figure 5B-D, Supplementary Figure 7C), either through activation or repression (defined in ^30^). We decided to focus on BLIMP1 for two reasons. First, its transcript is enriched in IL-10^+^ human B cells ^31^. Secondly, interrogation of a murine B cell ChIP- and RNA-seq data set ^32^ demonstrated that binding of BLIMP1 to the *IL10* locus correlates with increased IL-10 expression (Supplementary Figure 7D). Consistent with these data, we observed increased expression of BLIMP1 protein within the IL-10^+^ B cell populations (Figure 5E). To more thoroughly address its role, we performed gene targeting experiments on BLIMP1 in primary human B cells. TLR9 stimulation strongly upregulated BLIMP1 expression, whilst siRNA knockdown was able to reduce TLR9-dependent protein level expression by ∼60% (Figure 5F-H). In agreement with a central role in regulating IL-10, siRNA-mediated knockdown of BLIMP1 reduced IL-10 expression by 50-90%, whereas TNFα production was preserved (Figure 5I). Together, these data demonstrate that BLIMP1 regulates IL-10 in human B cells, and its expression is dependent on cholesterol metabolism for its provision of GGPP.

**Figure. 5.**
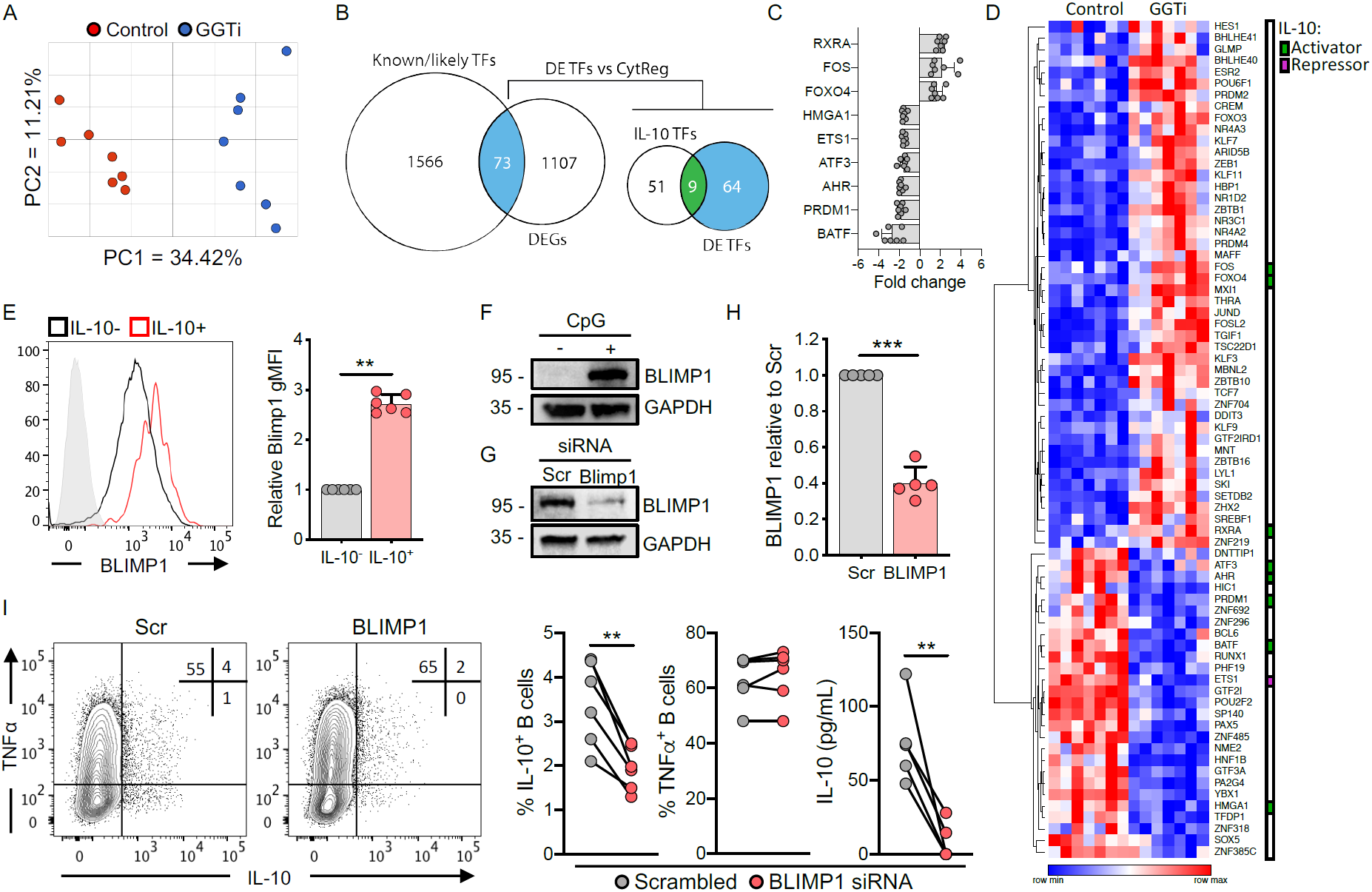
Geranylgeranyl pyrophosphate regulates IL-10 induction via BLIMP1. **A.** Principal component analysis of human B cells after stimulation through TLR9 ± geranylgeranyl transferase inhibition (GGTi), with analysis performed on total normalised counts generated from RNA-sequencing data, before statistical analysis (n=7). **B.** Workflow for the identification of putative IL-10 transcription factors. First, differentially expressed genes were cross referenced with known or likely human transcription factors, and the subsequently identified differentially expressed transcription factors were cross referenced with previously validated IL-10 transcription factors in either macrophages or T cells. Differentially regulated IL-10 transcription factors are shown in **(C)** and all differentially regulated transcription factors are shown in **(D). E.** Expression of BLIMP1 in human B cells within either IL-10^+^ or IL-10^−^ B cells after stimulation through TLR9. **F.** Western blot showing BLIMP1 expression in human B cells either in unstimulated or stimulated through TLR9 (CpG) after 40 hours (representative of three independent experiments). **G-H.** Western blot showing expression of BLIMP1 in human B cells after stimulation of TLR9 and nucleofection with either a scrambled control (Scr) or BLIMP1 siRNA. **I.** IL-10 and TNFα expression in human B cells stimulated through TLR9 after nucleofection with either Scr or BLIMP1 siRNA. Each data point represents individual donors. All data presented are mean ± SD where average values are shown. \*\**P*<0.01, \*\*\**P*<0.001 and all significant values are shown. Statistical analysis was conducted using a paired t-test

### Mevalonate kinase deficient patients generate poor regulatory B cell responses

Our data demonstrated that cholesterol metabolism is essential for B cell derived IL-10 production. We therefore sought to validate our findings in the context of human inflammatory disease. To address this, we studied MKD patients, who carry a partial loss-of-function mutation in the mevalonate kinase gene, whose cognate substrate lies directly downstream of HMG-CoA in the metabolic pathway (highlighted in Supplementary Figure 1A). These patients present with a complex and severe autoinflammatory syndrome ^25^. We first compared peripheral regulatory B cells from MKD patients to age and sex matched healthy controls (Supplementary Figure 8A). We observed similar frequencies of B10 cells, whereas CD24^hi^CD38^hi^ B cells were largely absent (Figure 6A-B). This is in line with a general shift from a naïve to memory like B cell phenotype in MKD patients (Supplementary Figure 8B). We next examined if dysregulated cholesterol metabolism in patients with MKD leads to an inability to mount an IL-10 response, which could contribute to disease persistence or exacerbation. In line with this hypothesis, MKD patients generated poor IL-10 responses after stimulation through TLR9 (30-70% reduction versus controls, Figure 6C), whilst TNFα expression was enhanced in 2 of 4 donors (Supplementary Figure 8C). Interestingly, the defect in IL-10 production could be reversed by the addition of GGPP, indicating that at least in part, this defect was due to a relative deficiency of the isoprenyl metabolite (Figure 6C).

**Figure. 6.**
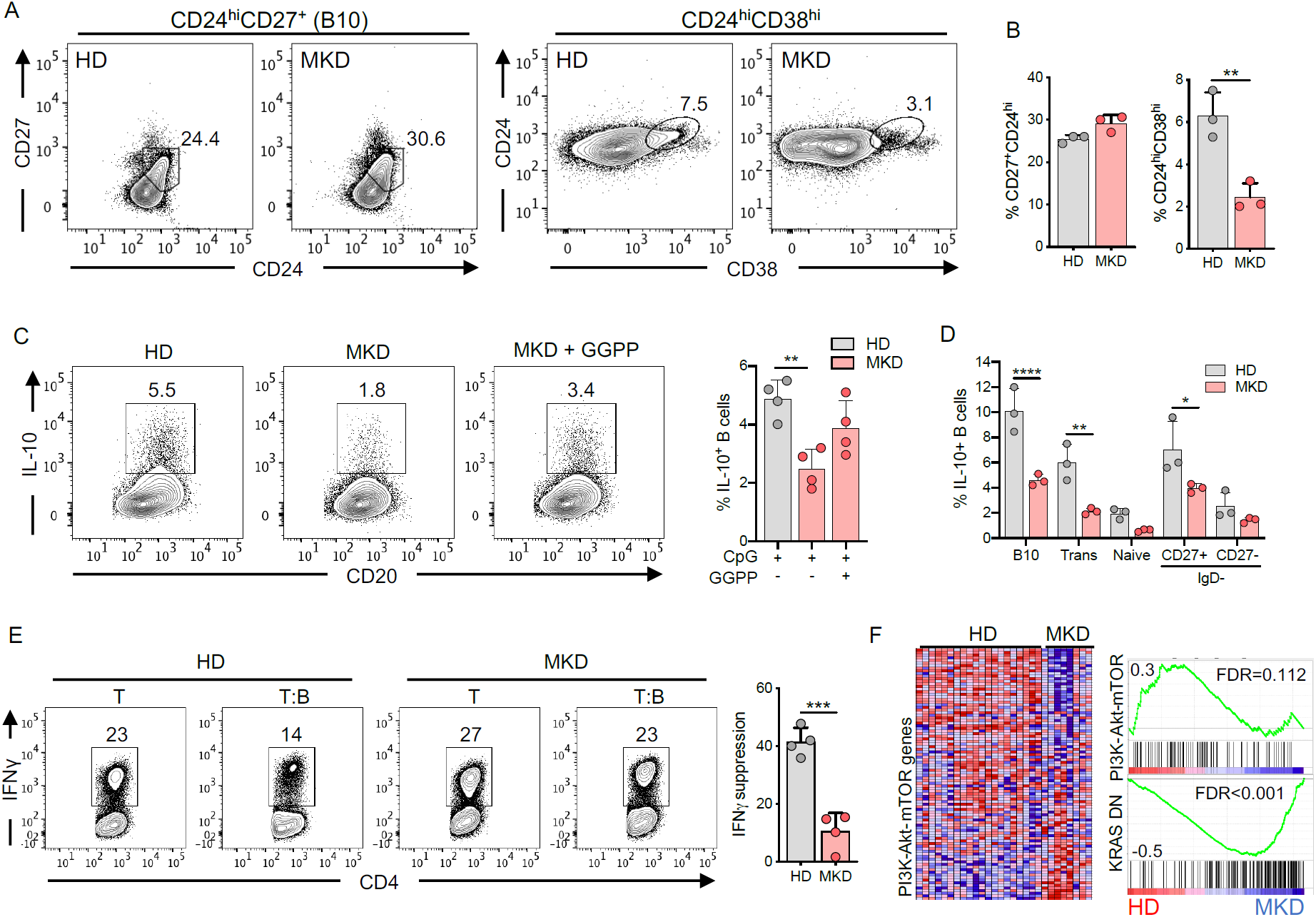
MKD patients generate a poor regulatory response across B cell populations, due to GGPP deficiency. **A-B.** Frequencies of B10 (CD24^hi^CD27^+^) and CD24^hi^CD38^hi^ regulatory B cells in mevalonate kinase deficient (MKD) patients, compared to age and sex matched healthy donors (HD). **C.** IL-10 expression in MKD patients and HD after stimulation through TLR9 ± geranylgeranyl pyrophosphate (GGPP). **D.** IL-10 expression within B10, CD24^hi^CD38^hi^, naïve, and memory populations, showing the differential contribution of B cell phenotypes to IL-10 production. **E.** IFNγ production in human CD4^+^ T cells of healthy controls and MKD patients after co-culture with autologous TLR9-activated B cells. Protocol is outlined in Fig. 1C. **F.** Gene set enrichment analysis conducted on gene expression profiles from an independent dataset (GSE43553) on PBMCs *ex vivo* in MKD patients (n=8) or HD (n=20). Heatmap shows all genes involved in ‘Hallmark’ PI3K-Akt-mTOR gene set collection from MSigDB, and their relative expression in MKD vs HD. Enrichment plots show PI3K-Akt-mTOR and KRAS DN (genes downregulated by KRAS activation) in MKD vs HD. Each data point represents individual donors. All data presented are mean +/-SD where average values are shown. \**P*<0.05, \*\**P*<0.01, \*\*\**P*<0.001, \*\*\*\**P*<0.0001 and all significant values are shown. Statistical analysis was conducted using Friedman’s test with Dunn’s multiple comparisons, or a paired t-test

As with our previous observations, IL-10 expression was reduced across all B cell phenotypes, including a ∼3-fold reduction within B10 and CD24^hi^CD38^hi^ B cells (Figure 6D, gating in Supplementary Figure 4B). This defect in IL-10 production was associated with functional impairment, as MKD B cells were unable to suppress IFNγ production by autologous CD4^+^ T cells, when compared to B cells from healthy controls (Figure 6E). Moreover, pre-treatment of MKD B cells with exogenous GGPP prior to co-culture, was able to restore regulatory capacity in 2 of 2 donors tested (Supplementary Figure 8D), suggesting that a GGPP deficiency impairs both IL-10 expression and defective suppressive function. To examine whether MKD imposed more a systemic defect on IL-10 producing immune cells we investigated whether T cell responses from MKD patients were also defective in IL-10 expression. In contrast to B cells, T cells were able to induce effective IL-10 responses, alongside comparable levels of other cytokines including TNFα, IFNγ, IL-2 and IL-17 (Supplementary Figure 8E).

Finally, to corroborate our results, we interrogated an independent dataset (GSE43553) that performed global gene expression analysis in *ex vivo* PBMCs from healthy controls (n=20) and MKD patients (n=8). Gene set enrichment analysis identified defective Ras (KRAS) and PI3K-Akt-mTOR signaling pathways in MKD patients when compared to healthy donors, in line with our own observations (Figure 6F). Collectively, and consistent with our *in vitro* findings, these data suggest that dysregulated cholesterol metabolism *in* vivo, as seen in MKD patients, results in impaired regulatory B cell responses.

## Discussion

This study provides the first description of the metabolic requirements for IL-10 producing B cells, arguing for a central reliance on cholesterol metabolism. Our data point to a model in which synthesis of GGPP prior to stimulation permits transduction of receptor signaling cascades necessary for IL-10 expression. As a consequence, the transcription factor BLIMP1 is induced, which then promotes IL-10 gene expression. We propose that this has direct relevance *in vivo*, as IL-10 producing B cells from patients who carry mutations in the cholesterol metabolic pathway phenocopy our *in vitro* findings.

Investigations into B cell metabolism have focused primarily on either antibody production or activation induced metabolic reprogramming ^18–20,33^. As alluded to above, there was no understanding of the metabolic requirements for regulatory B cells. Here we demonstrate that cholesterol metabolism is critical in mediating the regulatory capacity of human B cells through its control of IL-10. Interestingly, cholesterol metabolism has been implicated in regulatory T cell function through a mechanism dependent on ICOS and CTLA-4, whilst having no effect on IL-10 expression ^34^. Although the authors did not explore this, we anticipate that regulatory T cells may also possess significant GGPP dependency, as regulatory T cell function is especially reliant on PI3Kδ activity ^35^. Therefore, whilst cholesterol metabolism appears to regulate different effector molecules between the cell types, it may be that regulatory T and B cells rely more heavily on cholesterol metabolism due to the necessity for potent GGPP-dependent PI3Kδ activity.

Much of the experimental data regarding immunity and metabolism have suggested a paradigm in which the integration of metabolic pathways controlling cell fate arises as a direct consequence of immune cell activation. Based on the ability of cholesterol metabolism to control induction of a regulatory program in human B cells by modulating TLR9 signaling, we propose that the programming of immune responses through cholesterol metabolism may differ in this regard. We propose that isoprenyl modifications constitute a metabolic pre-programming event. This pre-programming requires the generation of GGPP prior to cellular stimulation (rather than upon or as a direct consequence of stimulation), that has the effect of fine-tuning signaling cascades. This notion of pre-programming is consistent with data showing the GGPP-dependent constitutive localization of Ras family proteins at the cell membrane and endosomes, and that blockade of cholesterol metabolism retains Ras in the cytoplasm ^36,37^. In other words, the state of signaling intermediates is preset by the state of cholesterol metabolism of the quiescent cell at any given point.

Our finding that either GGPP deficiency or inhibition of its cognate enzyme GGTase was sufficient to block IL-10 production suggested that both the metabolite and its enzyme are absolutely required for the induction of a regulatory phenotype. Notably, inhibition of FTase was unable to inhibit IL-10 production, likely due to the differing targets for geranylgeranylation and farnesylation ^27^. This suggests that cholesterol metabolism drives the anti-inflammatory function of B cells primarily through the synthesis of the isoprenyl group GGPP. These observations contrast with previous reports suggesting that isoprenylation largely mediates the restriction of pro-inflammatory cytokines, including TNFα, IL-6, and IL-1β. It should be noted however, that these data were derived from studies of murine macrophages and intestinal epithelial cells ^23,38,39^. Nonetheless, whilst human B cells are known to co-express inflammatory cytokines TNFα and IL-6 alongside IL-10 ^40^, we saw no alteration in their capability to produce these upon blockade of GGTase. This suggests that pro-inflammatory cytokine production relies less heavily on cholesterol metabolism. Our finding that cholesterol metabolism regulates PI3K signaling may somewhat explain this, given that PI3K can differentially regulate pro-versus anti-inflammatory cytokine production upon TLR ligation ^41–43^.

We found that the mechanistic control of cholesterol metabolism revolves around its ability to regulate PI3Kδ-AKT signaling downstream of TLR9. Although we documented expression of all isoforms of PI3K, we observed that PI3Kδ, to a greater extent than PI3Kα or PI3Kγ, contributed to IL-10 expression in human B cells upon TLR9 ligation. This finding is consistent with a recent observation identifying IL-10 producing B cells as the primary pathological cell type in a model of activated PI3Kδ syndrome, driving *Streptococcus pneumoniae* persistence ^14^. We further suggest that GSK3 inactivation downstream of GGPP-dependent PI3Kδ-AKT signaling is also required for optimal IL-10 expression. Consistent with this, AKT-driven phosphorylation of GSK3 on Ser9 is known to differentially regulate cytokine production in monocytes ^28^. Moreover, we observed that reduced IL-10 expression induced in the context of defective isoprenylation could be rescued through inhibition of GSK3. In line with our findings, GSK3 inhibition in both innate cells ^28^ and Th1/2/17 cells ^44^ potentiates IL-10 expression. This provides a common framework in all immune cells, whereby restriction of GSK3 activity upon receptor ligation is necessary for IL-10 expression.

Downstream of TLR9 ligation, BLIMP1 was shown to positively regulate IL-10 expression, suggesting its role as a transcriptional regulator of IL-10 in human B cells. Analysis of publicly available ChIP-seq datasets in both murine and human B cells are in line with this finding, suggesting that IL-10 is a significantly enriched target of BLIMP1 ^32,45^. This observation is reminiscent of recent work highlighting the role for BLIMP1-expressing regulatory plasmablasts and plasma cells *in vivo* ^3,4,46,47^, but also in humans *ex vivo* ^47^. Whilst this previous work provided a correlative link between BLIMP1 and IL-10, we suggest that BLIMP1 is a direct regulator of IL-10 in B cells. Therefore, it will be interesting in future work to further delineate a direct link and understand if there is a common mechanism regulating B cell derived IL-10 in both murine and human B cells. Additionally, several interesting candidates such as AHR and BATF were also identified in our screen, which warrant further investigation.

To validate our findings in the context of human disease, we investigated patients with dysregulated cholesterol metabolism. MKD patients carry a mutation in the mevalonate kinase gene. As a consequence, their ability to convert mevalonate to mevalonate-5-phosphate is severely impaired. In keeping with our findings following perturbation of cholesterol metabolism in healthy B cells *in vitro*, we observed poor regulatory responses in B cell from these patients. This anti-inflammatory defect uncovered an unappreciated dimension to the spectrum of MKD. This included a reduced ability to produce IL-10, associated with a functional impairment in restricting T cell responses. In both cases, supplementation of GGPP was able to reverse this defect, suggesting that a lack of metabolic flux through the pathway might be contributing. In agreement with this, MKD patients have been demonstrated to accumulate unprenylated Ras proteins ^48^. To date, the causative factor driving disease pathology has been defined as excessive inflammatory cytokine production, driven through increased macrophage driven IL-1β production ^23,49^, but also through the induction of a trained immunity phenotype driven by accumulated mevalonate ^50^. Here we also provide evidence that CD4^+^ T cells in MKD patients induce cytokine responses equivalent to those in healthy donors, suggesting this effect is intrinsic to B cells. Given the role of regulatory B cells in restricting cellular responses, we propose this B cell defect could contribute to the relapsing and remitting nature of the disease. Moreover, it has been shown that B cell derived IL-10 is important in T follicular helper organization, and restricting excessive antibody responses via interaction with Tfh cells ^51,52^. It is therefore tempting to speculate that these patients generate inappropriately inflammatory B cell responses in lymphoid organs. This would go some way to explain the clinical observations in MKD patients who are typically diagnosed in early childhood with high circulating IgD and IgA levels ^25^.

In summary, these results argue that cholesterol metabolism acts as a central metabolic pathway promoting an intrinsic anti-inflammatory program in B cells, driving IL-10 production and subsequent suppression and restriction of immune responses.

## Supplementary Figures

**Figure S1.**
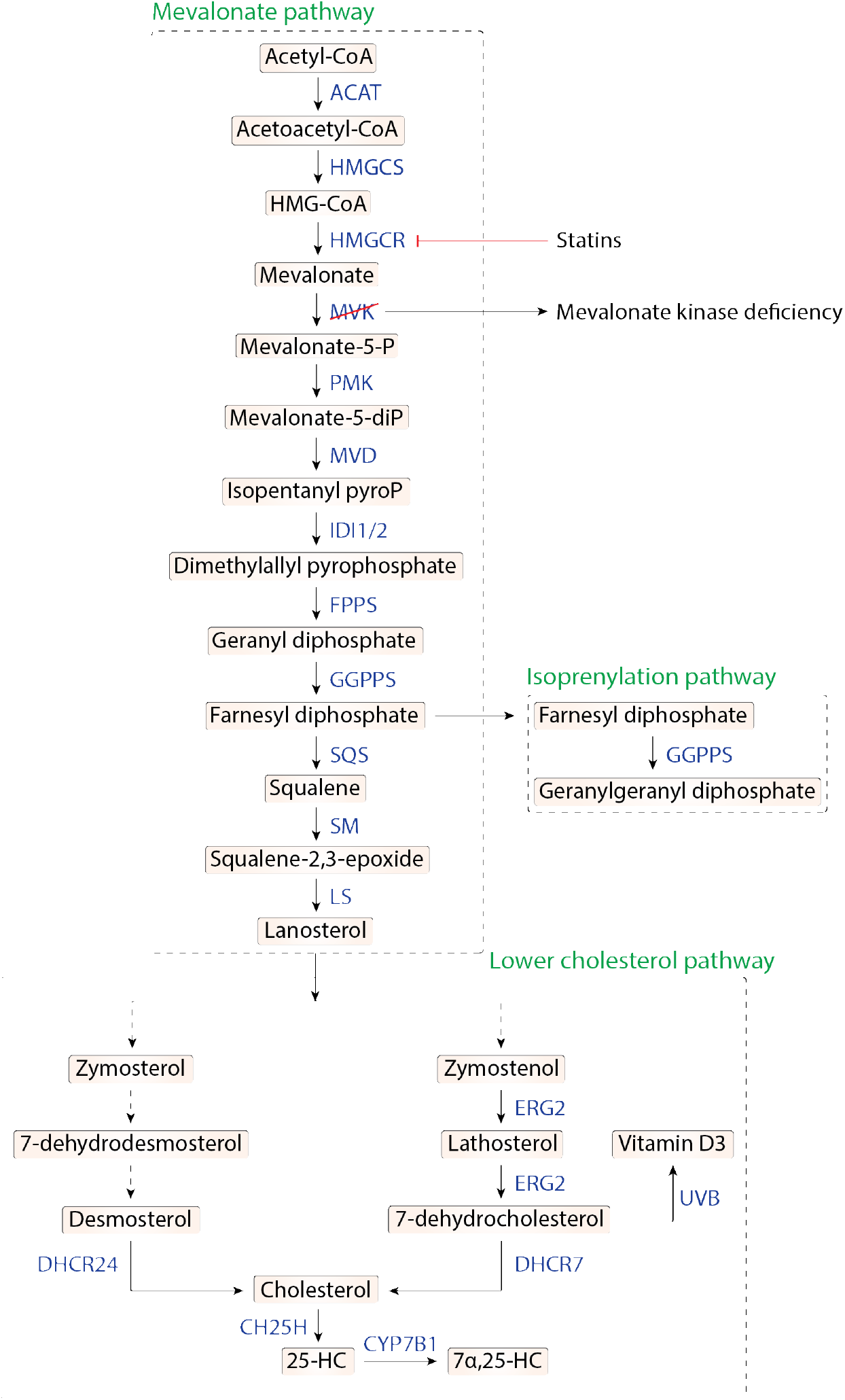
Outline of cholesterol metabolism showing key metabolites and enzyme of the pathway. An outline of key metabolites and enzymes in the multi-step conversion of acetyl-CoA to cholesterol and its derivatives, generally termed cholesterol metabolism. MKD patients suffer a mutation in mevalonate kinase (MVK) enzyme, and progress to a severe autoinflammatory syndrome.

**Figure S2.**
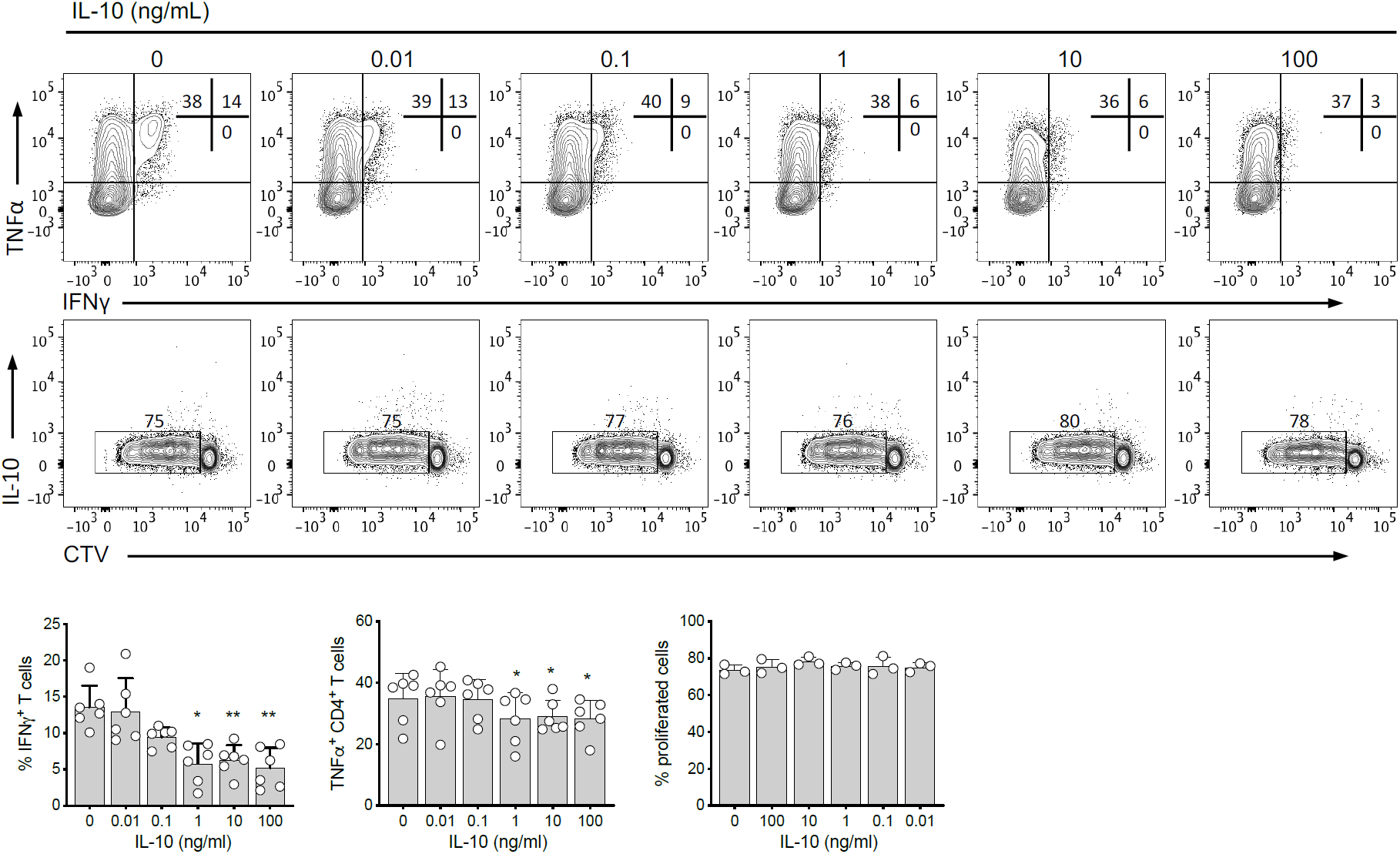
Direct effect of IL-10 on human CD4 T cell proliferation, TNFα, and IFNγ, cytokine production. Representative plots and quantification of IFNγ, TNFα expression, and proliferation in CD4^+^ T cells after stimulation with αCD3/28, titration of recombinant IL-10, and culture for 4 days. Each data point represents individual donors. All data presented are mean ± SD. \**P*<0.05, \*\**P*<0.01 and all significant values are shown. Statistical analysis was conducted using Friedman’s test with Dunn’s multiple comparisons

**Figure S3.**
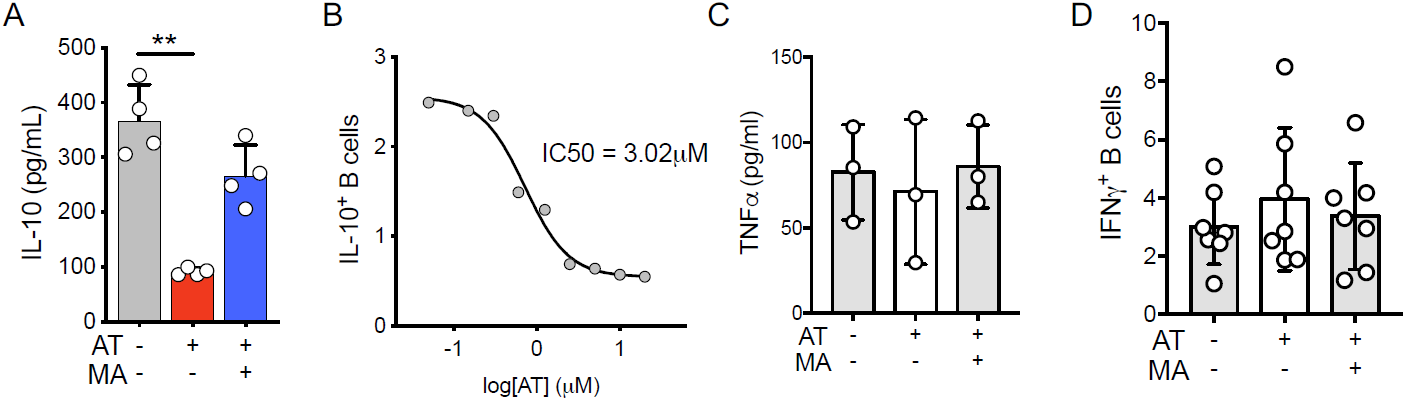
Cholesterol metabolism specifically regulates IL-10 expression. **A.** IL-10 secretion from human B cells after TLR9 stimulation, ± atorvastatin (AT) ± mevalonate (MA) (n=4). **B.** IC50 of IL-10 expression in human B cells stimulated through TLR9 in the presence of titrated levels of AT (n=4). **C.** Expression of TNFα (ELISA) and IFNγ (flow cytometry) in human B cells stimulated with TLR9 ± AT ± MA. Each data point represents individual donors. All data presented are mean ± SD. \*\**P*<0.01 and all significant values are shown. Statistical analysis was conducted using Friedman’s test with Dunn’s multiple comparisons

**Figure S4.**
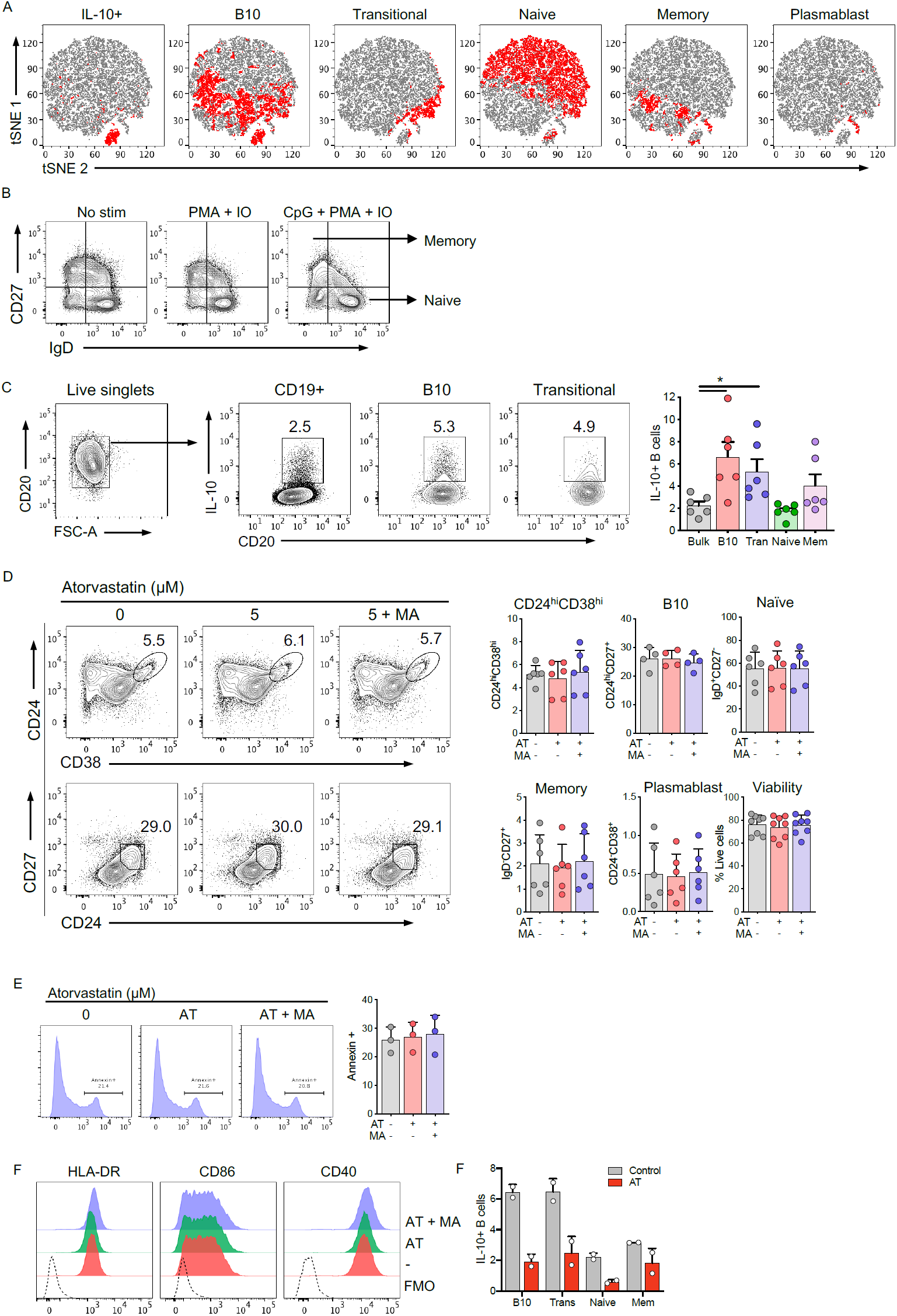
IL-10 expression within B cell populations, and the contribution of cholesterol metabolism to B cell phenotype. **A.** Representative tSNE analysis of human B cells stimulated through TLR9, highlighting specific populations (red) within total CD19^+^ B cells (grey) (n=3). Population gates are as follows: CD24^hi^CD27^+^ (B10) CD24^hi^CD38^hi^ (transitional), IgD^+^CD27^−^ (naïve), IgD^−^CD27^+^ (memory), CD24^−^CD38^+^CD27^+^ (plasmablast). **B.** Gating of naïve versus memory B cells after CpG stimulation. CpG stimulation results in the downregulation of CD27, in comparison to unstimulated cells, or phorbol 12-myristate 13-acetate and ionomycin stimulated cells. Memory cells were defined from the upper left quadrant, whereas naïve cells were defined in the lower right **C**. Percentage of IL-10^+^ human B cells within each population (bulk (CD19^+^), B10, transitional, naïve, and memory). **D-E.** Proportional analysis of B cell populations upon stimulation with TLR9 ± atorvastatin (AT) ± mevalonate (MA), and viability measured by live/dead stain or annexin positivity **(D). F.** Representative expression of HLA-DR, CD86, and CD40 on B cells after TLR9 stimulation ± AT ± MA. G. IL-10 expression within human B cell populations after stimulation through TLR9 ± AT. Each data point represents individual donors. All data presented are mean ± SD. \**P*<0.05 and all significant values are shown.

**Figure S5.**
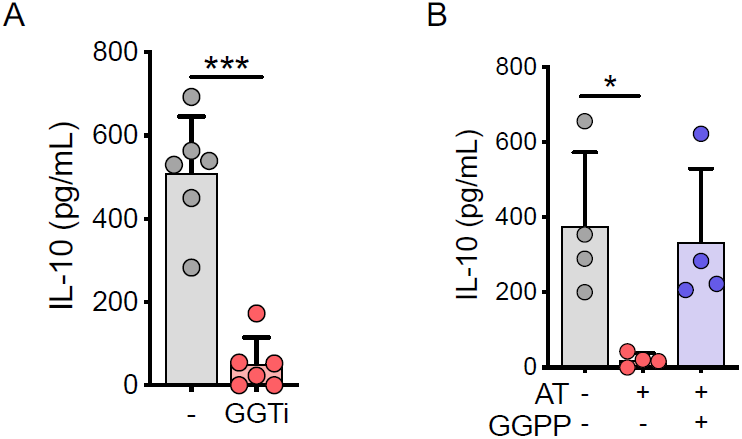
Cholesterol metabolism regulates IL-10 via GGTase and GGPP. **A.** IL-10 secretion in TLR9 stimulated human B cells in the presence of geranylgeranyl transferase inhibition (GGTi), measured by ELISA. **B.** IL-10 secretion in TLR9 stimulated human B cells ± atorvastatin (AT) ± geranylgeranyl pyrophosphate (GGPP). Each data point represents individual donors. All data presented are mean ± SD. \**P*<0.05, \*\*\**P*<0.001 and all significant values are shown. Statistical analysis was conducted using Friedman’s test with Dunn’s multiple comparisons, or a paired t-test

**Figure S6.**
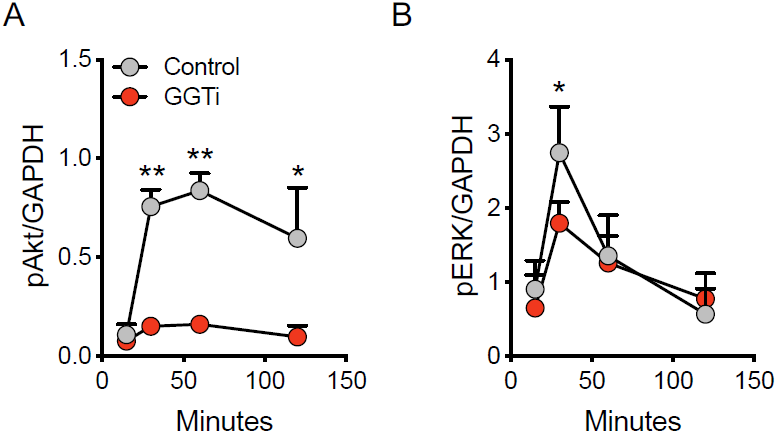
GGTase and GGPP regulate IL-10 induction downstream of TLR9 via PI3K, AKT, and ERK. **A-B.** Quantification of AKT **(A)** and ERK **(B)** phosphorylation over time in human B cells after stimulation through TLR9 ± geranylgeranyl transferase inhibitor (GGTi). All data presented are mean ± SD. \**P*<0.05, \*\**P*<0.01 and all significant values are shown. Statistical analysis was conducted using Friedman’s test with Dunn’s multiple comparisons

**Figure S7.**
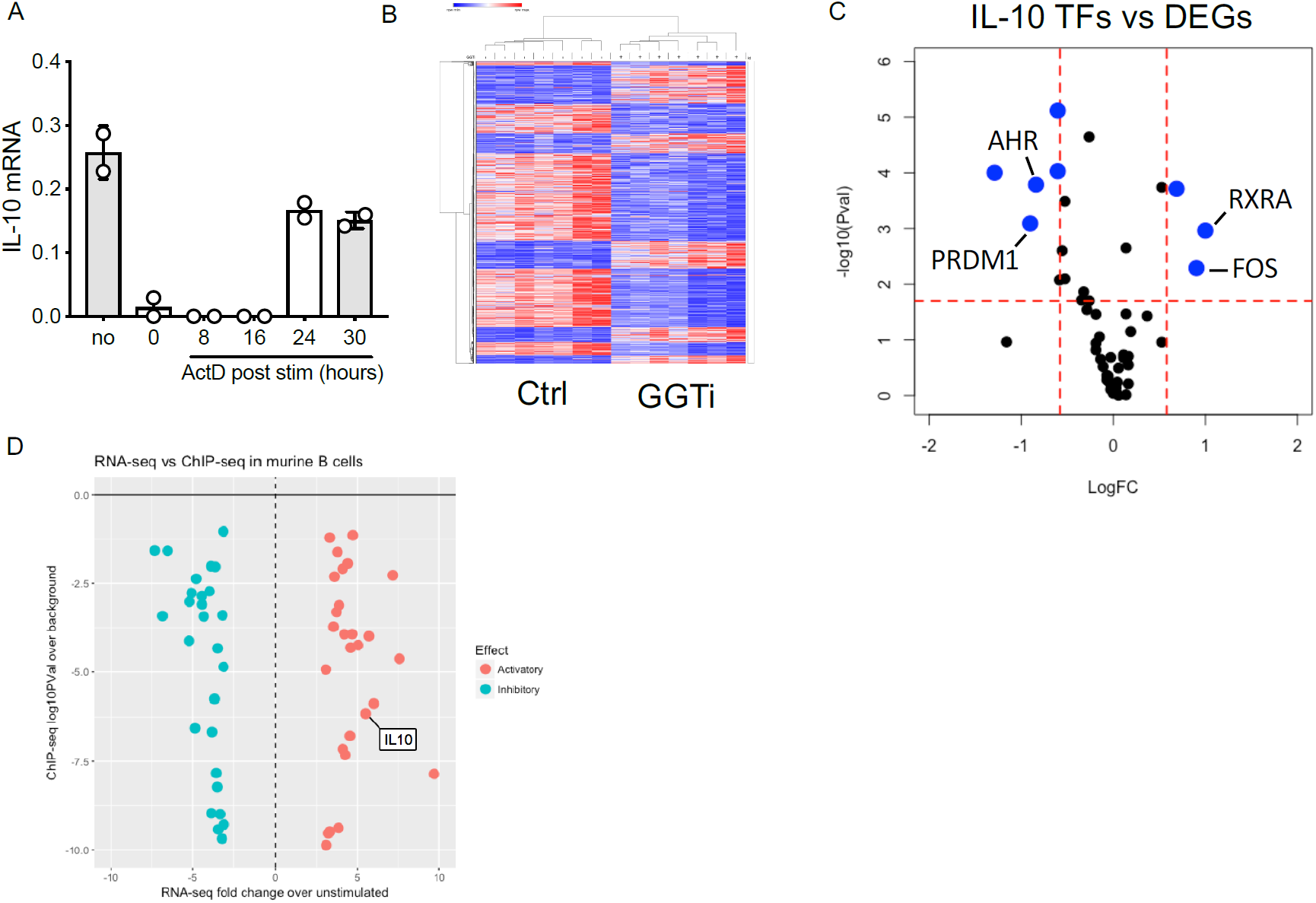
A transcriptional event mediated by GGTase after TLR9 stimulation is required for the expression of a transcription factor necessary for IL-10. **A.** qRT-PCR analysis of IL-10 expression in B cells following TLR9 stimulation ± actinomysin D (ActD) added at the indicated time post-stimulation, with all cells acquired at 40 hours. **B.** A heatmap representing the global profile of differentially expressed genes (FDR<0.05, Fold change >1.5) in human B cells following stimulation through TLR9 ± geranylgeranyl transferase inhibition (GGTi). **C.** All previously experimentally validated IL-10 transcription factors, and their FDR and fold change in our data set. Red dashed lines represent FDR=0.05 and fold change=1.5. **D.** ChIP-seq (y-axis) vs RNA-seq (x-axis) data from GSE71698, showing enriched targets of BLIMP1, and the correlative change in gene expression. Each data point represents individual donors. All data presented are mean ± SD.

**Figure S8.**
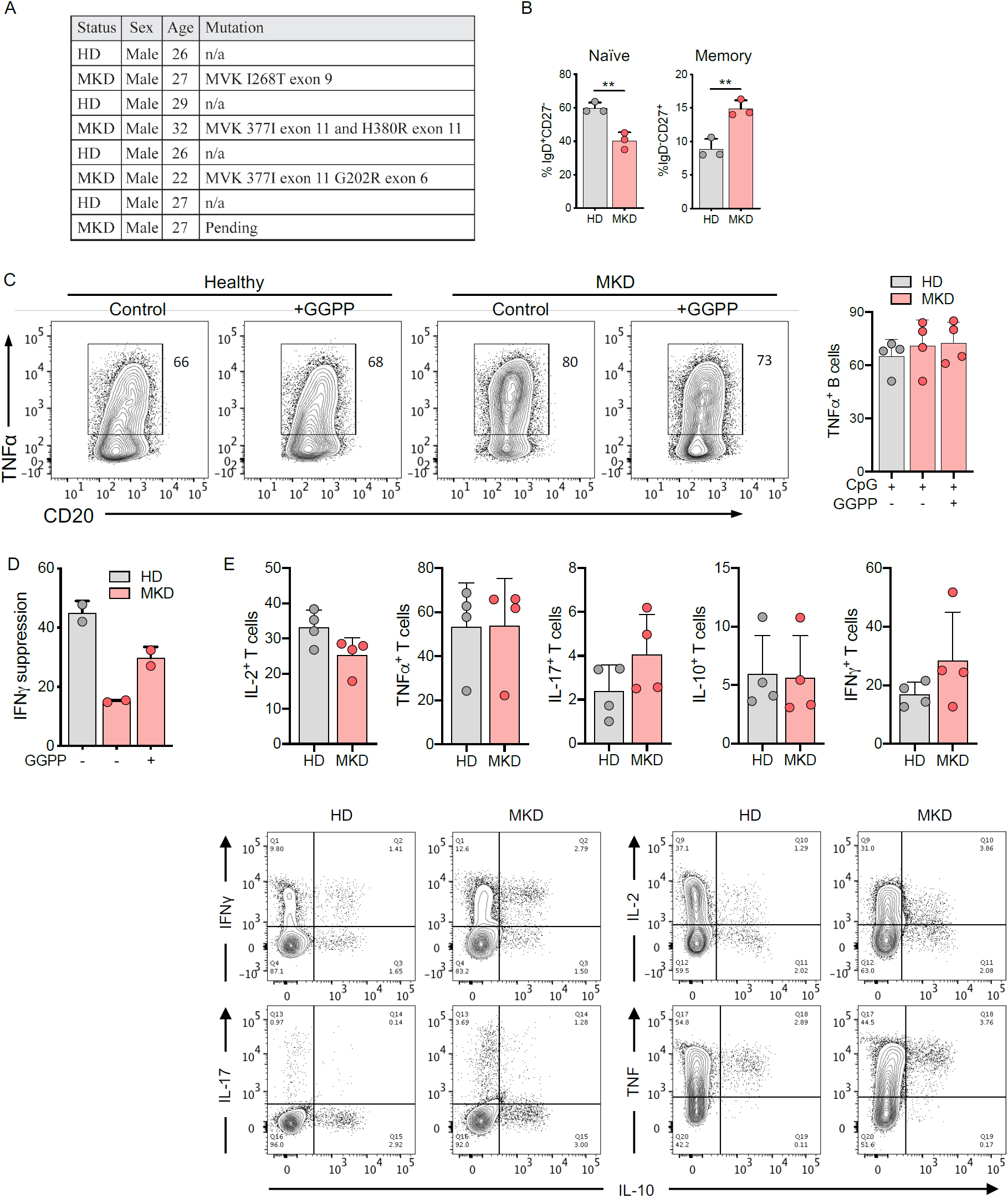
MKD patients show poor regulatory B cell responses, but normal T cell responses. **A.** Information for healthy donors (HD) and mevalonate kinase deficient (MKD) patients used in this study. **B. N**aive versus memory B cell populations in MKD patients relative to healthy donors, as a percentage of total CD19^+^ B cells. **C.** TNFα expression in human B cells from HD or MKD patients stimulated through TLR9 ± geranylgeranyl pyrophosphate (GGPP). **D.** IFNγ production in human CD4^+^ T cells of healthy controls and MKD patients after co-culture with autologous TLR9 activated B cells ± GGPP treated B cells prior to co-culture. **E**. Cytokine production in CD4^+^ T cells from MKD patients and healthy controls. Each data point represents individual donors. All data presented are mean ± SD. All significant values are shown. Statistical analysis was conducted using Friedman’s test with Dunn’s multiple comparisons

## Acknowledgements

We thank all the healthy donors and patients for their support. We also thank the Genomics Centre at King’s College London. This work was supported by the IMI-funded project BeTheCure, grant number 115142-2 and from the EU/EFPIA Innovative Medicines Initiative 2 Joint Undertaking RTCure grant no. 777357 (E.P. and A.P.C.); the National Institute for Health Research Biomedical Research Centre, at Guy’s and St Thomas’ NHS Foundation Trust and King’s College London (J.A.B.)

## Author contributions

J.A.B. designed, performed, and interpreted experiments, and wrote the manuscript, H.A.P. contributed to experimental design, performed experiments, and contributed to manuscript preparation, T.H. performed experiments, A.C. and K.O. contributed to experimental design and provided mice, M.W. recruited patients and extracted patient samples, H.J.L contributed to experimental design and provided patient samples, C.K. supervised research, and contributed to experimental design, A.P.C. and E.P. conceived the project, designed experiments, supervised research, and contributed to manuscript preparation.

## Declaration of interests

The authors declare no competing interests

## Methods

### Cell isolation and culture

Blood samples were obtained locally from healthy donors, and mevalonate kinase deficient patients were recruited from Royal Free Hospital NHS Foundation Trust. Human: Peripheral blood mononuclear cells were isolated by density gradient centrifugation using Lymphoprep (Alere Technologies). Fresh CD19+ B cells and CD4+ T cells were subsequently isolated by magnetic cell sorting, based on CD19 and CD4 positive or CD4 negative selection (MACS, Miltenyi Biotech); purity was consistently >97%. Cells were cultured in RPMI-1640 (Sigma) with 2mM L-glutamine (Sigma), 0.1g/L sodium bicarbonate (Sigma), supplemented with 100U/mL penicillin and 0.1mg/mL Streptomycin (PAA), and 10mM Hepes (PAA), in round bottom 96-well Nunclon plates (Thermo Scientific) at 0.3 × 106 cells/well. B cells were activated with TLR9 ligand CpG ODN 2006 (1μM, Invivogen), alongside IL-2 (25U/ml, Proleukin, Novartis) typically for 40 hours, or as indicated. All inhibitors and metabolites were extensively titrated prior to use, and reagent details are as follows: MK-2206 (Akt, 100nM), FTi (Farnesyl transferase, 5μM), GGTi-298 (Geranylgeranyl transferase, 5μM), CHIR-99021 (GSK3α/β, 5μM), atorvastatin (HMG-CoA reductase, 5μM), U0126 (MEK, 1μM), HS-173 (p110α, 1pM-1μM), IP-549 (p110α, 1pM-1μM), nemiralisib (p110δ, 1pM-1μM), (R)-mevalonic acid (250nM), and geranylgeranyl pyrophosphate (2μM). Mouse: Total splenocytes were isolated from 8-15-week-old mice, red blood cells were lysed (Biolegend), and resulting single cell suspensions were cultured in the above media in flat bottom 48-well Nunclon plates (ThermoFisher), at 1× 10^6^ cells/well. Wild type, p110δ E1020K and p110δ D910A mice used in the study have been described previously (24). B cells were stimulated with CpG ODN 1826 (1μM, Invivogen) in the presence of IL-2 (25U/ml) for 48 hours.

### T and B cell co-culture

Autologous CD4^+^ T and CD19^+^ B cells were isolated from healthy donors or MKD patients as described above. B cells were incubated overnight with CpG (1μM), IL-2 (25U/ml), +/- atorvastatin, +/- mevalonic acid. In parallel, CD4^+^ T cells were stimulated with plate bound αCD3 and αCD28 (both 1μg/ml, Biolegend). After 12 hours, B cells were washed twice, added to the T cells at a 1:1 ratio, and cultured for 4 days +/- αIL10 antibody +/- isotype control (5μg/ml, both Biolegend). Assessment of the direct effect of IL-10 on human CD4 T cells was performed by plate bound αCD3/28 (1μg/ml) stimulated, cell trace violet labelled T cells, alongside incubation of the indicated concentration of IL-10 for 4 days.

### Flow Cytometry

Antibodies used were as follows: IL-10-PE (JES3-19F1), TNFα-BV421 (Mab11), IFNγ-PE/Cy7 (4S.B3), CD20-AF647 (2H7), CD24-BV605 (ML5), CD27-FITC (M-T271), CD38-PE/Cy7 (HB-7), IgD-BV421 (IA6-2), CD4-FITC (A161A1), HLA-DR-PerCP/Cy5.5 (L243), CD86-BV421 (IT2.2), CD40-PE (5C3): all Biolegend, and pGSK3ser9-PE (REA436, Miltenyi). For intracellular cytokine detection, cells were restimulated in the final 3 (human) or 5 (mice) hours of culture with phorbol 12-myristate 13-acetate (50□g/mL) and ionomyocin (500ng/mL, Sigma) in the presence of Brefeldin A, and GolgiStop (both 1μl/ml, both BD Biosciences). For surface staining, cells were harvested and incubated with Fixable Viability Dye eFluor780 (eBiosciences) for 15 minutes in phosphate buffered saline (PBS), followed by the appropriate volume of antibody diluted in 0.5% bovine serum albumin (BSA) in PBS for 20 minutes, all at 4°C. Cells were then washed and fixed in 3% paraformaldehyde (Electron Microscopy Sciences) for 15 minutes at room temperature. For intracellular cytokine staining, cells were incubated with the appropriate volume of antibody, diluted in 0.1% saponin in 0.5% BSA in PBS for 45 minutes at room temperature or 4°C overnight. For staining of transcription factors, or phospho-flow, FoxP3/Transcription factor buffer set was used, as per manufacturer’s instructions (eBiosciences). Staining using a secondary antibody against the primary was undertaken either at room temperature for 1 hour, or 4°C overnight. Cells were acquired using a BD LSRFortessa or FACSCanto II (BD Biosciences), and analysis conducted using FlowJo V.10.1 software (Tree Star Inc.).

### tSNE analysis

Samples were stained using fluorophore-conjugated antibodies (as above) against: CD20, CD24, CD27, CD38, IgD, and IL-10. tSNE analysis was conducted in FlowJo (V.10.1). After exclusion of dead cells, doublets, and cell debris, downsampling was conducted to generate 30,000 events per sample. The parameters used for tSNE analysis were: iterations = 1000, perplexity = 30, learning rate = 700.

### RNA extraction and qRT-PCR analysis

Harvested cells were lysed in TRIzol (Ambion, Life Technologies), and stored at −20°C until RNA extraction by phenol-chloroform phase separation. Briefly, RNA was eluted by chloroform, and subsequently precipitated using ice cold isopropanol for 1 hour at −20°C. If necessary, RNA clean-up was conducted by reprecipitation of RNA in 0.5 volumes of 7.5M ammonium acetate, 2.5 volumes 100% ice cold ethanol, and 1μl GlycoBlue overnight at −20°C. 40ng of RNA was added for reverse transcription, using qPCRBIO cDNA Synthesis Kit (PCR Biosystems) as per manufacturer’s instructions. qPCR was conducted using 7900HT Fast Real-Time PCR System (ThermoFisher Scientific). Primers used were: IL10 (Hs00961622_m1), IL6 (Hs00174131_m1), or LTA (Hs04188773_g1), multiplexed with a VIC-labelled 18S probe (all Thermofisher Scientific) in PCR Biosystems mastermix.

### ELISA

Sandwich ELISA was used to detect supernatant analytes. All cell free supernatants harvested at indicated time points were stored at −20°C until analysis. ELISAs for IL-10, IFNγ (both R&D systems), and TNFα (both Biolegend) were conducted according to manufacturer’s protocols and detected on a Victor 1420 multilabel counter (Perkin Elmer) quantifying concentrations drawn from a standard curve on each plate.

### Western Blotting

Antibodies used were as follows: BLIMP1 (646702, R&D Systems), pAktser473 (D9E), pERKThr202/Tyr204 (D12.14.4E), and GAPDH (14C10, all Cell Signaling Technologies). After culture, cells were harvested, washed in ice cold PBS, and lysates immediately extracted, or pellets were snap frozen on dry ice for 1 minute and kept at −80°C until lysis. Whole cell lysates were extracted using RIPA Buffer (Cell Signaling Technology) with 1X Protease inhibitor cocktail. Lysates were added to 4X Laemmli Sample Buffer (Bio Rad) with 10% 2-mercaptoethanol, and resolved on SDS-PAGE gels, transferred to PVDF membranes, blocked (Tris, 5% BSA, 0.05% Tween20) and probed with the desired antibody. Immunoblots were developed with anti-rabbit-HRP (Dako) secondary antibody, and proteins visualized by SuperSignal chemiluminescent reaction (Pierce) in a ChemiDoc station (BioRad).

### siRNA knockdown

siRNA knockdown was conducted by Amaxa nucleofection using the nucleofector 2b machine, as per manufacturers protocol. siRNA knockdown was optimized, resulting in the used of 2-3 million cells per condition, and 500nM of either BLIMP1 targeted siRNA (Silencer Select assay ID s1992) or scrambled control (Silencer Select Negative control, both from ThermoFisher Sceintific). Cells were electroporated using program U-015, resuspended in warm fully supplemented culture medium, and left to recover for 15 minutes before stimulation. Knockdown efficiency was determined by western blot at 40 hours post stimulation.

### RNA sequencing

CD19^+^ B cells were stimulated with CpG (1μM) and IL-2 (25U/ml) +/- GGTi-298 for 12 hours. RNA was isolated by column centrifugation using RNeasy Plus mini kit (with gDNA removal step) as per manufacturer’s instructions (Qiagen). Library preparation was completed using NEBNext Ultra Directional RNA Library Prep Kit for Illumina. Depletion of ribosomal RNA was performed using Next rRNA Depletion kit (New England BioLabs). RNA quality was confirmed by bioanalyser (Agilent 2100 Bioanalyzer G2938B), resulting in a mean RIN score of 8.1, ranging from 7.3-8.7. Paired-end sequencing was then conducted using the HiSeq 2500 platform (Illumina). Raw data was checked for quality using FASTQC. Processing of the raw data involving alignment (STAR) and annotation (hg19) were done using Partek. After annotation, reads per million normalized data were then used for statistical analysis. Inclusion criteria for significantly differentially expressed genes was a false discovery rate of <0.05 and a fold change of greater than 1.5x. Subsequent processing and visualization of the data was completed in RStudio or Morpheus (Broad Institute, Boston, MA).

### Gene set enrichment analysis

Gene set enrichment analysis used on dataset GSE43553 was conducted with GSEA v4.0.2 (Broad Institute). Gene expression data was used from 20 healthy controls (GSM1065214 to GSM1065233) and 8 MKD patients (GSM1065234 to GSM1065241). Gene sets were compared to the gene set database h.all.v7.0.symbols.gmt [Hallmarks], using 1000 permutations, the ‘weighted’ enrichment statistic, and ‘Signal2Noise’ for gene ranking.

### Statistics

All statistical analysis was conducted using GraphPad Prism v7.0 (GraphPad, San Diego, CA, USA). Paired data was analysed through either two-way ANOVA with Dunnett’s test for multiple comparisons, or Freidman’s test with Dunn’s test for multiple comparisons. Unpaired data was assessed through ANOVA with Tukey’s test for multiple comparisons. Finally, two-group comparison was performed either through unpaired or paired T test. A P value of < 0.05 was considered statistically significant, and all statistically significant values are shown.

### Study approval

Blood samples were obtained locally from healthy donors, and mevalonate kinase deficient patients were recruited from Royal Free Hospital NHS Foundation Trust. Written informed consent was received from all healthy donors and patients used in this study, approved by the Bromley Research Ethics Committee (REC06/Q0705/20). Animal experiments were performed according to the Animals (Scientific Procedures) Act 1986, license PPL 70/7661 and approved by the Babraham Institute Animal Welfare and Ethics Review Body.

